# neurolib: a simulation framework for whole-brain neural mass modeling

**DOI:** 10.1101/2021.02.18.431886

**Authors:** Caglar Cakan, Nikola Jajcay, Klaus Obermayer

## Abstract

*neurolib* is a computational framework for whole-brain modeling written in Python. It provides a set of neural mass models that represent the average activity of a brain region on a mesoscopic scale. In a whole-brain network model, brain regions are connected with each other based on biologically informed structural connectivity, i.e. the connectome of the brain. *neurolib* can load structural and functional datasets, set up a whole-brain model, manage its parameters, simulate it, and organize its outputs for later analysis. The activity of each brain region can be converted into a simulated BOLD signal in order to calibrate the model against empirical data from functional magnetic resonance imaging (fMRI). Extensive model analysis is possible using a parameter exploration module, which allows one to characterize the model’s behavior given a set of changing parameters. An optimization module can fit a model to multimodal empirical data using an evolutionary algorithm. *neurolib* is designed to be extendable such that custom neural mass models can be implemented easily, offering a versatile platform for computational neuroscientists for prototyping models, managing large numerical experiments, studying the structure-function relationship of brain networks, and for performing in-silico optimization of whole-brain models.

## Introduction

Mathematical modeling and computer simulations are fundamental for understanding complex natural systems. This is especially true in the field of computational neuroscience, where models are used to represent neural systems at many different scales. At the macroscopic scale, we can study whole-brain networks that model a brain that consists of brain regions which are coupled via long-range axonal connections. A number of technological and theoretical advancements have transformed the concept of whole-brain modeling from an experimental proof-of-concept into a widely used method that is part of the toolkit of today’s computational neuroscientists, the first of which is the widespread availability of computational resources.

An integral part of this development can be attributed to the success of mathematical neural mass models representing the population activity of a neural network, often by using mean-field theory^1,2^ which employs methods from statistical physics^3^. While microscopic simulations of neural systems often rely on large spiking neural network simulations where the membrane voltage of every neuron is simulated and kept track of, neural mass models typically consist of a system of differential equations that govern the macroscopic variables of a large system, such as the population firing rate. Therefore, these models are considered useful for representing the average activity of a large neural population, e.g., a brain area. Biophysically realistic population models are often derived from networks of excitatory (E) and inhibitory (I) spiking neurons by assuming the number of neurons to be very large, their connectivity sparse and random, and the post-synaptic currents to be small^4^. At the other end of the spectrum of neural mass models, simple phenomenological oscillator models^5-8^ are also used to represent the activity of a single brain area, sacrificing biophysical realism for computational and analytical simplicity.

In the past, whole-brain models have been employed in a wide range of problems, including demonstrating the ability of whole-brain models to reproduce BOLD correlations from functional magnetic resonance imaging (fMRI) during resting-state^9,10^ and sleep^11^, explaining features of EEG^12^ and MEG^5,7^ recordings, studying the role of signal transmission delays between brain areas^13,14^, the differential effects of neuromodulators^8,15^, modeling electrical stimulation of the brain in-silico^16-18^, or explaining the propagation of brain waves^19^ such as in slow-wave sleep^20^. Previous work often focused on finding the parameters of optimal working points of a whole-brain model, given a functional dataset^21^.

However, although it is clear that whole-brain modeling has become a widely-used paradigm in computational neuroscience, many researchers rely on a custom code base for their simulation pipeline. This can result in slow performance, avoidable work due to repetitive implementations, the use of lengthy boilerplate code, and, more generally, a state in which the reproduction of scientific results is hard. In order to address these points, we present *neurolib*, a computational framework and a Python library, which helps users to set up whole-brain simulations. With *neurolib*, parameter explorations of models can be conducted in large-scale parallel simulations. *neurolib* also offers an optimization module for fitting models to experimental data, such as from fMRI or EEG, using evolutionary algorithms. Custom neural mass models can be implemented easily into the existing code base. The main goal of *eurolib* is to provide a fast and reliable framework for numerical experiments that encourages customization, depending on the individual needs of the researcher. *neurolib* is available as free open source software released under the MIT license.

## Results

### Whole-brain modeling

A whole-brain model is a network model which consists of coupled brain regions (see Fig. 1). Each brain region is represented by a neural mass model which is connected to other brain regions according to the underlying network structure of the brain, also known as the connectome^22^. The structural connectivity of the brain is typically measured using diffusion tensor imaging (DTI) which is used to infer the long-range axonal white matter tracts in the brain, a method known as fiber tractography. When combined with a parcellation scheme that divides the brain into different brain regions, also known as an atlas, the brain can be represented as a brain network with the brain regions being its nodes and the white matter tracts being its edges. Figure 1 shows structural connectivity matrices that represent the number of fibers and the average fiber length between any two regions. In a simulation, each brain area produces activity, for example a population firing rate and a BOLD signal. To assess the validity of a model, the simulated output can then be compared to empirical brain recordings.

**Figure 1.**
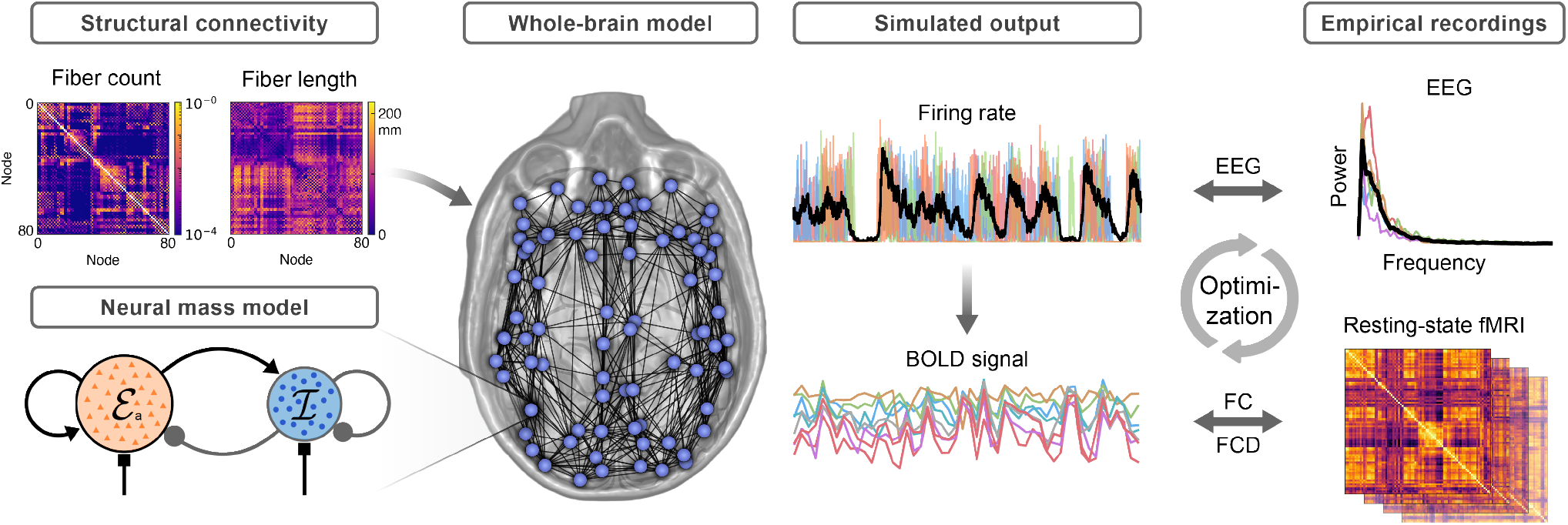
Construction of a whole-brain model. Structural connectivity from DTI tractography is combined with a neural mass model that represents a single brain area. The depicted neural mass model consists of an excitatory (red) and an inhibitory (blue) subpopulation. The default output of each region, e.g. the excitatory firing rate, is converted to a BOLD signal using the hemodynamic Balloon-Windkessel model. For model optimization, the models’ output is compared to empirical data, such as an EEG power spectrum or to fMRI functional connectivity (FC) and its temporal dynamics (FCD).

### Framework architecture

In the following, we will describe the design principles of *neurolib* and provide a brief summary of the structure of the Python package here (see Fig. 2). Later, the individual parts of the framework will be discussed in more detail. At the core of *neurolib* is the `Model` base class from which all models inherit their functionality. The base class initializes and runs models, manages parameters, and handles simulation outputs. To reduce memory footprint of long simulations, chunkwise integration can be performed using the autochunk feature. The outputs of a model can be converted into a BOLD signal using a hemodynamic model which allows for a comparison of the simulated outputs to empirical fMRI data. The `Dataset` class handles structural and functional datasets. A set of post-processing functions and a `Signal` class is provided for computing functional connectivity (FC) matrices, applying temporal filters to model outputs, computing power spectra, and more. The simulation pipeline interacts with two additional modules that provide parameter exploration capabilities using the BoxSearch class, and enable model optimization using the `Evolution` class. Both modules utilize the `ParameterSpace` class which provides the appropriate parameters values for exploration and optimization.

**Figure 2.**
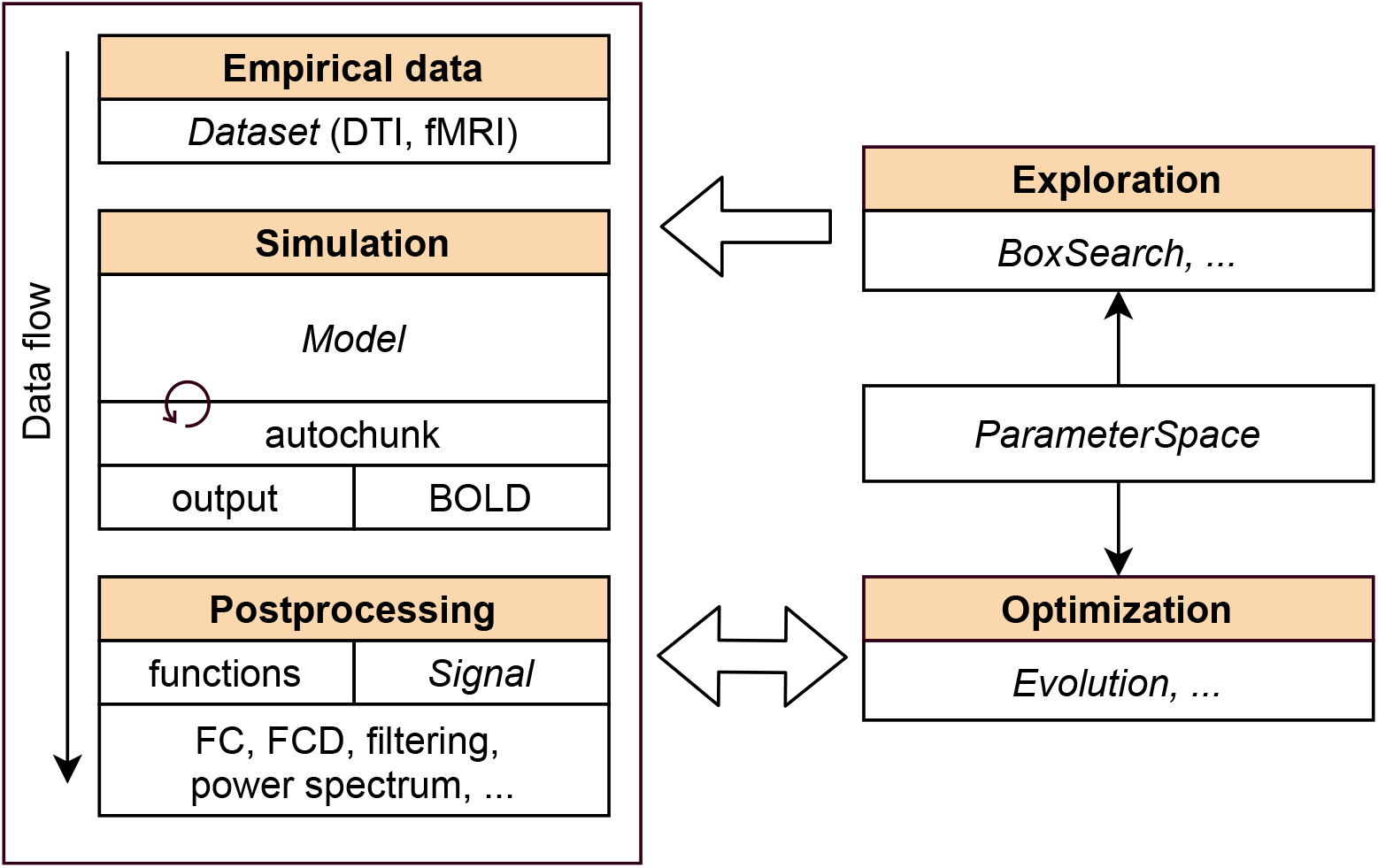
Framework architecture. Class names are in cursive letters.

### Neural mass models

Several neural mass models for simulating the activity of a brain area are implemented in *neurolib* (see Table 1). Some neural mass models, for example the ALN model^23,24^ or the Wilson-Cowan model^25,26^, consist of multiple neural populations, namely an excitatory (E) and an inhibitory (I) one, which are referred to as subpopulations in order to distinguish it from an entire brain area, which we refer to as a node. Every brain area is a node coupled to other nodes in the whole-brain network. It should be noted that some phenomenological models like the Hopf model^27^ only have a single variable that represents neural activity, and, therefore, the distinction between E and I subpopulations does not apply.

**Table 1.**
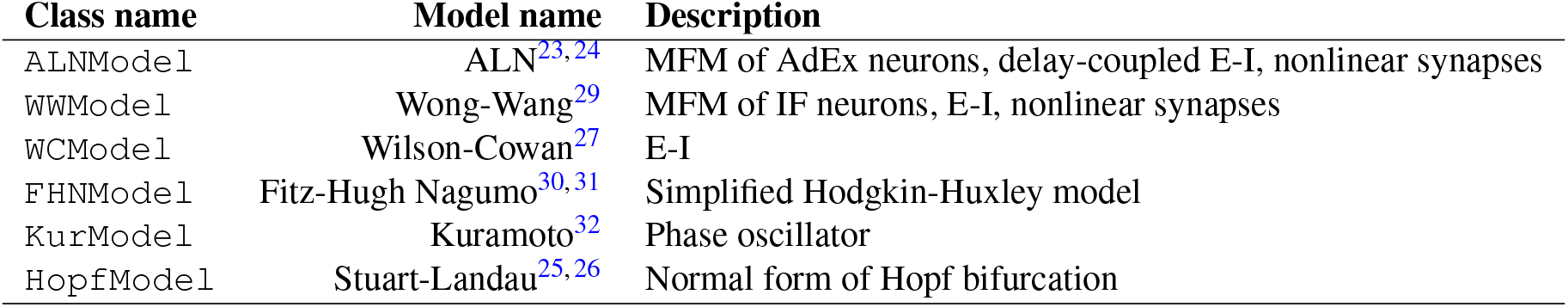
Implemented neural mass models. Mean-field models of spiking neurons are abbreviated as MFM. Models with excitatory and inhibitory subpopulations are abbreviated as E-I. The integrate-and-fire neuron model is abbreviated as IF and the adaptive exponential integrate-and-fire model as AdEx.

| Class name | Model name | Description |
| --- | --- | --- |
| ALNModel | ALN<sup>23,24</sup> | MFM of AdEx neurons, delay-coupled E-I, nonlinear synapses |
| WWModel | Wong-Wang<sup>29</sup> | MFM of IF neurons, E-I, nonlinear synapses |
| WCModel | Wilson-Cowan<sup>27</sup> | E-I |
| FHNModel | Fitz-Hugh Nagumo<sup>30,31</sup> | Simplified Hodgkin-Huxley model |
| KurModel | Kuramoto<sup>32</sup> | Phase oscillator |
| HopfModel | Stuart-Landau<sup>25,26</sup> | Normal form of Hopf bifurcation |

### Phenomenological and biophysical models

Biophysically grounded neural mass models that are derived from an underlying network of spiking neurons produce an output that is a firing rate, akin to the mean spiking rate of a neural network. An example of such a model is the ALN model, which is based on a network of spiking adaptive exponential (AdEx) integrate-and-fire neurons^28^. Phenomenological models usually represent a simplified dynamical landscape of a neural network and produce outputs that are abstract and do not have physical units. An example is the Hopf model, where the system dynamics can be used to describe the transition from steady-state firing to neural oscillations^7^. The Wilson-Cowan model can be mentioned as a middle ground between simple and realistic. It describes the activity of excitatory and inhibitory neurons while relying on simplifications such as representing the fraction of active neurons, rather than the actual firing rate of the population, and uses an analytical firing rate transfer function.

### Model equations

The core module consists of a `Model` class that manages the whole-brain model and its parameters. Every model is implemented as a separate class that simulates the activity of the brain network. Typically, a model is implemented as a system of ordinary differential equations which can be expressed as

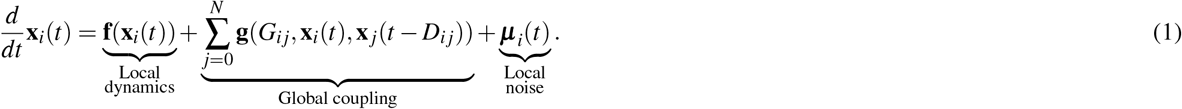

Here, the vector **x**_*i*_ = (*x*_*i*1_,…,*x_id_*) describes the *d*-dimensional state of the *i*-th brain region which follows the local node dynamics **f**, with *i* ∈ [0, *N* - 1] and *N* being the number of brain regions. This vector contains all state variables of the system, including for example, firing rates, and synaptic currents. The second term describes the coupling between the *i*-th and *j*-th brain regions, given by a coupling scheme **g**. This coupling term typically depends on the *N* × *N* adjacency matrix **G** (with elements *G_ij_*), the current state vector of the *i*-th brain area, and the time-delayed state vector of the *j*-th brain area, which itself depends on the *N* × *N* inter-areal signal delay matrix **D** (with elements *D_ij_*). The matrices **G** and **D** are defined by the empirical structural connectivity datasets. The third term *μ_i_*(*t*) represents a noise input to every node which is simulated as a stochastic process for each brain area (or subpopulation) individually.

The main difference between the implemented neural mass models is the local node dynamics **f**. Some models, such as the Hopf model, additionally support either an additive coupling scheme **g**, where the coupling term only depends on the activity of the afferent node ***x**_j_*, or a diffusive coupling scheme, where the coupling term depends on the difference between the activity of the afferent and efferent nodes, i.e. ***x**_i_* - ***x**_j_*. Other coupling schemes, such as non-linear coupling, can be also implemented by the user.

### Noise input

Every subpopulation α, with for example α ∈ {*E, I*}, of each brain area *i* (index omitted) receives an independent noise input *μ_α_*(*t*), which, for the models in Table 1, comes from an Ornstein-Uhlenbeck process^33^,

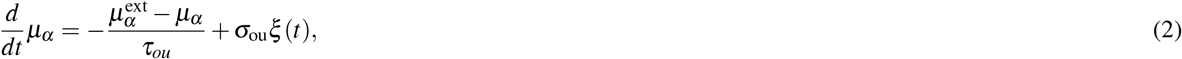

where 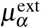 represents the mean of the process and can be thought of as a constant external input, *τ_ou_* is the time scale, and ξ(*t*) is a white noise process sampled from a normal distribution with zero mean and unit variance. The noise strength parameter σ_ou_ determines the standard deviation of the process and, therefore, the amplitude of fluctuations around the mean.

### Bifurcation diagrams

Figure 3 shows the bifurcation diagrams (or state spaces) of a single node of a selection of models in order of increasing complexity. The diagrams serve as a demonstration of the parameter exploration module of *neurolib*, which we will describe in more detail in a later section. Understanding the state space of a single node allows one to interpret its behavior in the coupled case. Starting from the Hopf model (Fig. 3 a), we can see how the transition to the oscillatory state is caused by an eponymous supercritical Hopf bifurcation controlled by the parameter *a*. Figures 3 b and 3 c show the time series of the activity variable (including noise) in the steady-state, i.e. a fixed point, for *a* < 0, and an oscillatory state, i.e. a limit cycle, for *a >* 0, respectively. The Wilson-Cowan model (Fig. 3 d) has two Hopf bifurcations, where the low-activity down-state is separated from the high-activity up-state by a limit cycle region in which the activity alternates between the E and I subpopulations. The time series of the activity variable in Fig. 3 e shows the system in the down-state with short excursions into the limit cycle due to the noise in the system, whereas in Fig. 3 f, the system is placed inside the limit cycle and reaches the up- and the down-state occasionally. The bifurcation diagram of the ALN model in Fig. 3 g has a more complex structure and its validity has been verified using large spiking network simulations before^24^. Here, we can see that the down-state and the up-state are separated by a limit cycle as well. Therefore, the bifurcation structure of the Wilson-Cowan model can be thought of as a simplified version of a slice going through the limit cycle of this two-dimensional diagram in the horizontal plane. In Fig. 3 h, the ALN model is placed in the down-state close to the limit cycle and the time series of the excitatory firing rate *r_e_* shows brief excursions into the oscillatory state. Without the adaptation mechanism that is derived from the underlying AdEx neuron, we can also observe a bistable regime in the bifurcation diagram Fig. 3 g, where both up- and down-state coexist. An example time series is shown in Fig. 3 i, where the activity shows noise-induced transitions between the up- and down-state. When the adaptation mechanism of the underlying AdEx model is enabled (not shown), the bistable region transforms into a new, slowly oscillating limit cycle^24,34^.

**Figure 3.**
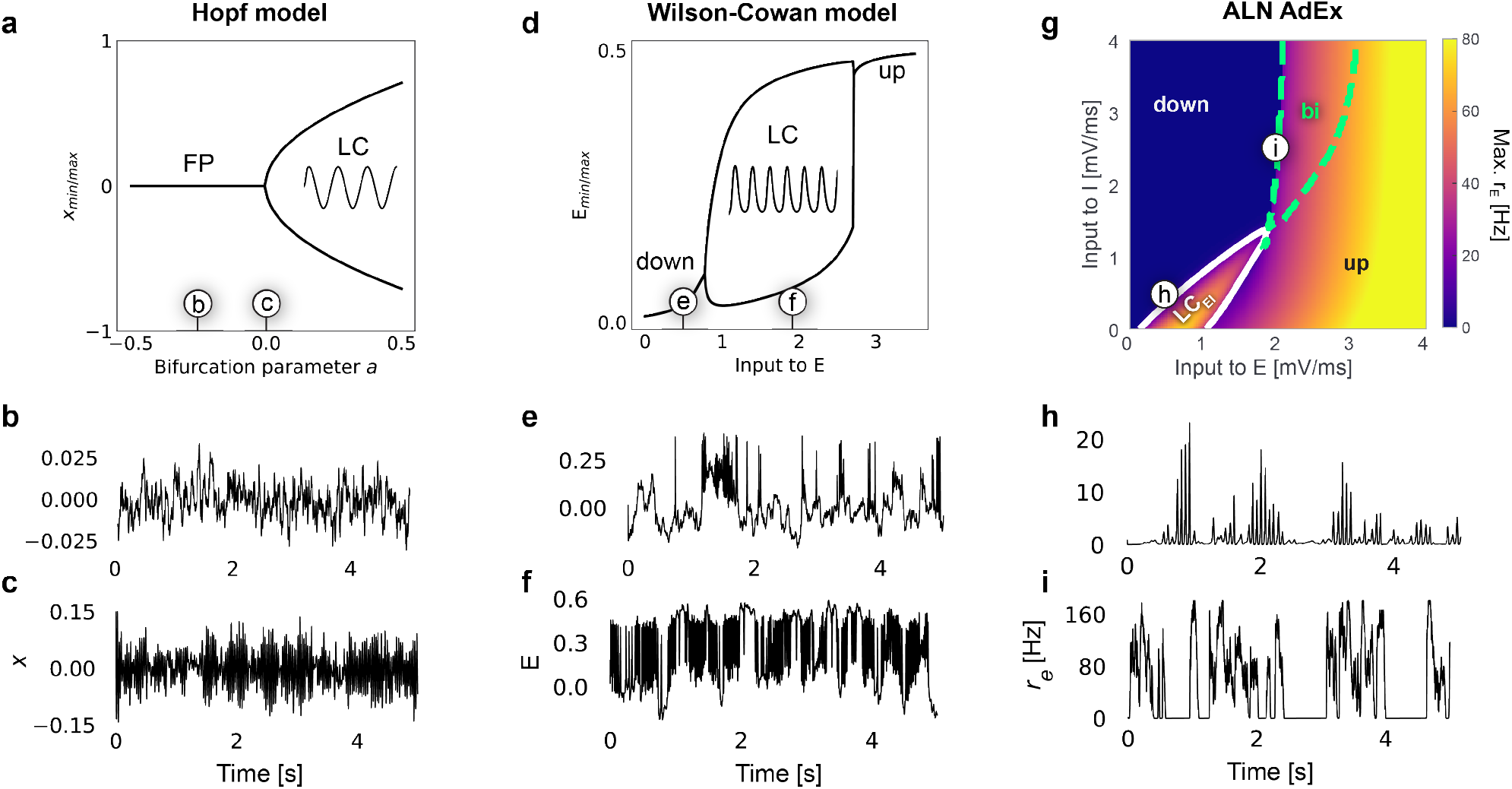
Overview of neural mass models. **(a)** Bifurcation diagram of the Hopf model. For *a* < 0, solutions converge to a fixed point (FP). For *a* > 0, solutions converge to a limit cycle (LC). **(b)** Example time series of *x* of the Hopf model in the FP with *a* = −0.25, noise strength σ_ou_ = 0.001, and noise time scale τ_ou_ = 20.0ms. **(c)** Time series at the bifurcation point *a* = 0 with the same noise properties as in (b). **(d)** Wilson-Cowan model with the activity of the excitatory population (E) plotted against the external input E_ext_. The system has two fixed-points with low (down) and high (up) activity and a limit cycle in between. **(e)** Time series of E with E_ext_ = 0.5, σ_ou_ = 0.01, and τ_ou_ = 100.0ms. **(f)** E_ext_ = 1.9. **(g)** Two-dimensional bifurcation diagram of the ALN model depending on the input currents to the excitatory (E) and inhibitory (I) population. The color denotes the maximum firing rate r_E_ of the E population. Relong-enough simulation time of 10 minutes in order tolong-enough simulation time of 10 minutes in order tolong-enough simulation time of 10 minutes in order togions of low-activity *down-states* (down) and high-activity *up-states* (up) are indicated. Dashed green contours indicate bistable (bi) regions where both states coexist. Solid white contour indicates oscillatory states within fast E-I limit cycle (LC_EI_). **(h)** Time series of the excitatory firing rate *r_e_* of the ALN model with mean input currents 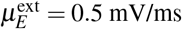 and 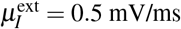 and noise parameters like in (e). **(i)** 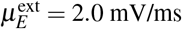 and 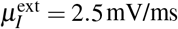 and σ_ou_ = 0.3. Each point in the bifurcation diagrams was simulated without noise for 5 seconds and the activity of the last second was used to determine the depicted state. All remaining parameters are at their default values.

### Numerical integration

The models are integrated using the Euler—Maruyama integration scheme^35^. The numerical integration is written explicitly in Python and then accelerated using the just-in-time compiler *numba*^36^, providing a performance similar to running native C code. Compared to pure Python code, this offers a speedup in simulation time on the order of 10^4^. For computational efficiency, each neural mass model is implemented as a coupled network such that the single-node case is a special case of a network with *N* = 1 nodes. Likewise, the noise process in Eq. 2 is also implemented within every model’s integration and then added to the appropriate state variables of the system, i.e., the membrane currents of E and I in the case of the ALN model.

### Example: Single node simulation

In the example in Listing 1, we load a single isolated (E-I) node of the `ALNModel`. The first step is to initialize the model. Every model has a set of default parameters which we can change by setting entries of the `params` dictionary attribute. To demonstrate this, we set the external noise strength which is simulated as an Ornstein-Uhlenbeck process with a standard deviation σ_ou_ = 0.1 (Eq. 2) and then run the model. The results from this simulation can be accessed via the `model` object’s attributes t which contains the simulation time steps, and `output` which contains the firing rate of the excitatory population. All other state variables of a model can be accessed via the dictionary attribute `outputs`. An example time series of the excitatory firing rate is shown in Figure 3 h.

**Listing 1.**
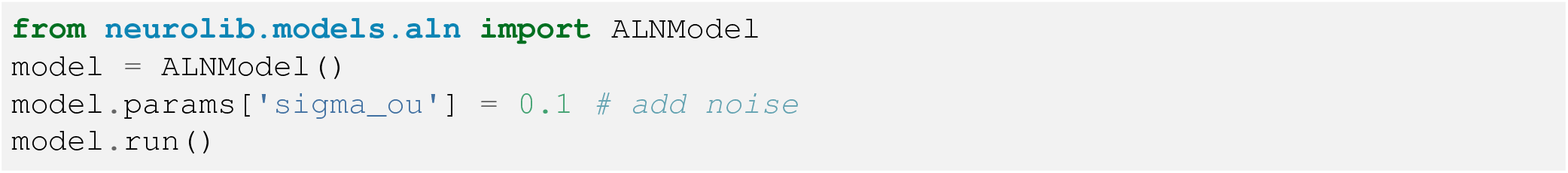
Single node simulation. A single node of the ALN mean-field model is simulated with noise.

### Empirical datasets

Empirical datasets are stored in the `data` directory. In each dataset, subject-wise functional and structural data are stored as MATLAB `.mat` matrices that can be opened in Python using SciPy’s `loadmat` function^37^. Structural data are *N* × *N*, and functional time series are *N* × *t* matrices, *N* being the number of brain regions and *t* the number of time steps. Example datasets are included in *neurolib* and custom datasets can be added by placing them in the dataset directory. Throughout this paper, we use preprocessed data from the ConnectomeDB of the Human Connectome Project (HCP)^38^. The handling of subject-wise datasets is done by the `Dataset` class, and the attributes in the following refer to its instances.

#### Structural DTI data

To simulate a whole-brain network model, first we need to load the structural connectivity matrices from a DTI dataset. The matrices are usually a result of processing DTI data and performing fiber tractography using software like *FSL*^39^ or *DSIStudio*^40^. For a given parcellation of the brain into *N* brain regions, these matrices are the *N* × *N* adjacency matrix `Cmat`, i.e. the structural connectivity matrix, which determines the coupling strengths between brain areas, and the fiber length matrix `Dmat` which determines the signal transmission delays. Example structural matrices are shown in Fig. 1. The datasets currently included in *neurolib* use the 80 cortical regions of the AAL2 atlas^41^ to define the brain areas and are sorted in a LRLR-ordering.

#### Connectivity matrix normalization

The elements of the structural connectivity matrix `Cmat` are typically the number of reconstructed fibers from DTI tractography. Since the number of fibers depends on the method and the parameters of the (probabilistic or deterministic) tractography, they need to be normalized using one of the three implemented methods. The first method `max` is to simply divide the entries of Cmat by the largest entry, such that the largest entry becomes 1. The second method `waytotal` divides the entries of each column of Cmat by the number fiber tracts generated from the respective brain region during probabilistic tractography in FSL, which is contained in the `waytotal.txt` file. The third method nvoxel divides the entries of each column of `Cmat` by the size, i.e., the number of voxels of the corresponding brain area. The last two methods yield asymmetric connectivity matrices, while the first one keeps them symmetric. All normalization steps are done on the subject-wise matrices (attributes `Cmats` and `Dmats`). Finally, group-averaged matrices are computed for the dataset and made available as the attributes `Cmat` and `Dmat`.

#### Functional MRI data

Subject-wise fMRI time series must be in a (*N* × *t*)-dimensional format, where *N* is the number of brain regions and *t* the length of the time series. Each region-wise time series represents the BOLD activity averaged across all voxels of that region, which can be also obtained from software like *FSL*. Functional connectivity (FC) captures the spatial correlation structure of the BOLD time series averaged across the entire time of the recording. Subject-wise FC matrices are accessible via the attribute FCs and are generated by computing the Pearson correlation of the time series between all regions, yielding a *N*× *N* matrix for each subject. Example FC matrices from resting-state fMRI recordings are shown in Fig. 1.

To capture the temporal fluctuations of time-dependent FC(t), which are lost when averaging across the entire recording time, functional connectivity dynamics matrices (`FCDs`) are computed as the element-wise Pearson correlation of time-dependent FC(t) matrices in a moving window across the BOLD time series of a chosen window length of, for example, 1 min. This yields a *t_FCD_ × t_FCD_* FCD matrix for each subject, with *t_FCD_* being the number of steps the window was moved.

### Example: Whole-brain simulation

In a whole-brain model, the main activity variables of each neural mass are coupled with each other. If the neural mass model has E and I subpopulations, the coupling is usually implemented between the activity variables of the E subpopulations, resulting in a whole-brain model with global excitation and local inhibition. The adjacency matrix `Cmat` determines the relative coupling strengths between all brain areas. The relative coupling strengths are multiplied by the global coupling strength parameter `Ke_gl` to determine the absolute coupling strengths. The elements of the delay matrix `Dmat` contain the average fiber lengths between any two brain regions in units of mm. The fiber length is divided by the signal transmission speed parameter `signalV` to determine the time delay for signal transmission. Given these matrices, we can initialize a brain network model by passing the data to the model’s constructor (Listing 2). We set a long-enough simulation time of 10 minutes in order to simulate the same length of the BOLD data as in the empirical dataset. Finally, we can choose to additionally simulate a BOLD signal by using the appropriate argument in the `run` method.

**Listing 2.**
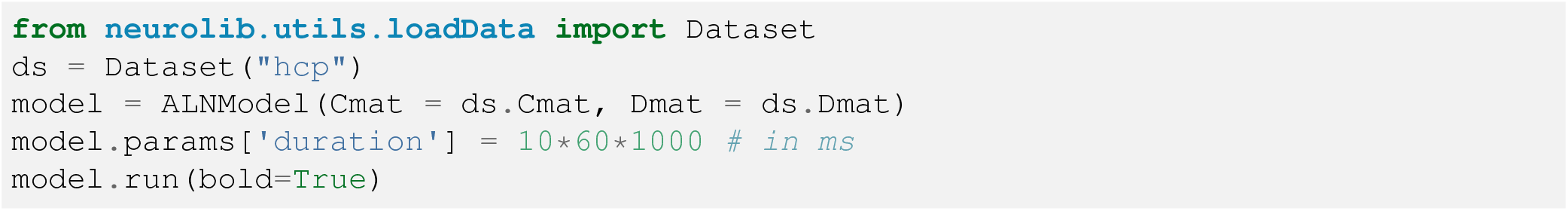
Brain network simulation. Structural connectivity and delay matrices are loaded from the HCP dataset and a whole-brain model is initialized. A simulation time of 10 minutes is set and the model is simulated with BOLD output.

#### BOLD model

Every brain area has a predefined default output variable which is one of its state variables. The default output variable of the ALN model, for example, is the firing rate of the excitatory subpopulation of every brain area. The default output can be used to simulate a BOLD signal using the implementation of the Balloon-Windkessel model^42-44^. The BOLD signal is governed by a set of differential equations that model the hemodynamic response of a brain area to neural activity. After integration, the BOLD signal is then subsampled at 0.5 Hz to match the sampling rate of fMRI recordings. The BOLD signal is integrated alongside the neural mass model and stored in the model’s outputs. To enable the simulation of the BOLD signal, the user passes the argument `bold = True` to the run method (see Listing 2).

### Example: Custom model implementation

In the following, we present how a custom model can be implemented in *neurolib*. Every model consists of two parts (see Listing 3). The first part is the class that implements the model and that inherits most of its functionality from the `Model` base class. The second part is the `timeIntegration()` function. In this example, we implement a linear model with the following equation

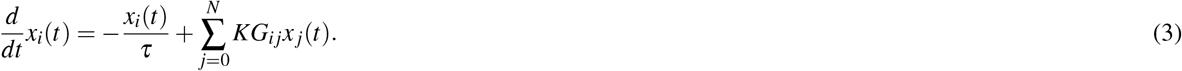

As before, this equation represents *N* nodes that are coupled in a network. *x_i_* are the elements of an *N*-dimensional state vector **x**, *τ* is the decay time constant, *G_ij_* are elements of the adjacency matrix **G**, and *K* is the global coupling strength. We implement this model as the class `LinearModel` in Listing 3.

In the definition of the model class, we specified some necessary information, such as the names of the state variables `state_vars`, the default output of the model `default_output`, and the variable names `init_vars`, holding the initial conditions at *t* = 0. The `timeIntegration()` function has two parts: One, in which the variables for the simulation are prepared, and another, where the actual time integration takes place, i.e., `njit_int()`. The latter has a decorator **@numba.njit** which ensures that the integration will be accelerated with the just-in-time compiler *numba.* The equations of the model are then integrated using the Euler integration scheme. This simple model can be run like the other models before, supports features like chunkwise integration (see below), and can produce a BOLD signal.

### Chunkwise integration for memory-intensive experiments

Some of the important applications of whole-brain modeling require very long simulation times in order to extract meaningful data from the model and to compare it to empirical recordings. Examples are computing BOLD correlations, such as FC and FCD matrices, from data with a very low sampling rate of around 0.5 Hz, or the computation of EEG power spectra over a long time period, or measuring event statistics based on, for example, transitions between up- and down-states^20^. This poses a major resource problem, since the neural dynamics is usually simulated with an integration time step of the order 0.1 ms or less, producing large amounts of data that an ordinary computer is not able to handle efficiently in its memory (RAM).

To overcome this issue, we designed a chunkwise integration scheme called *autochunk* which can be enabled by running a model using the command `model. run(chunkwise=True)`. It supports all models that follow the implementation guidelines. In this scheme, all dynamical equations are integrated for a short duration T_chunk_ (e.g. 10 seconds) as defined by the number of time steps of a chunk, `chunksize`. The chunk duration T_chun_k largely determines the amount of necessary RAM for a simulation and is typically a lot smaller than the total duration of the simulation T_total_. This means that the entire simulation will be integrated in ⌈*T*_total_/*T*_chunk_⌉ chunks.

**Listing 3.**
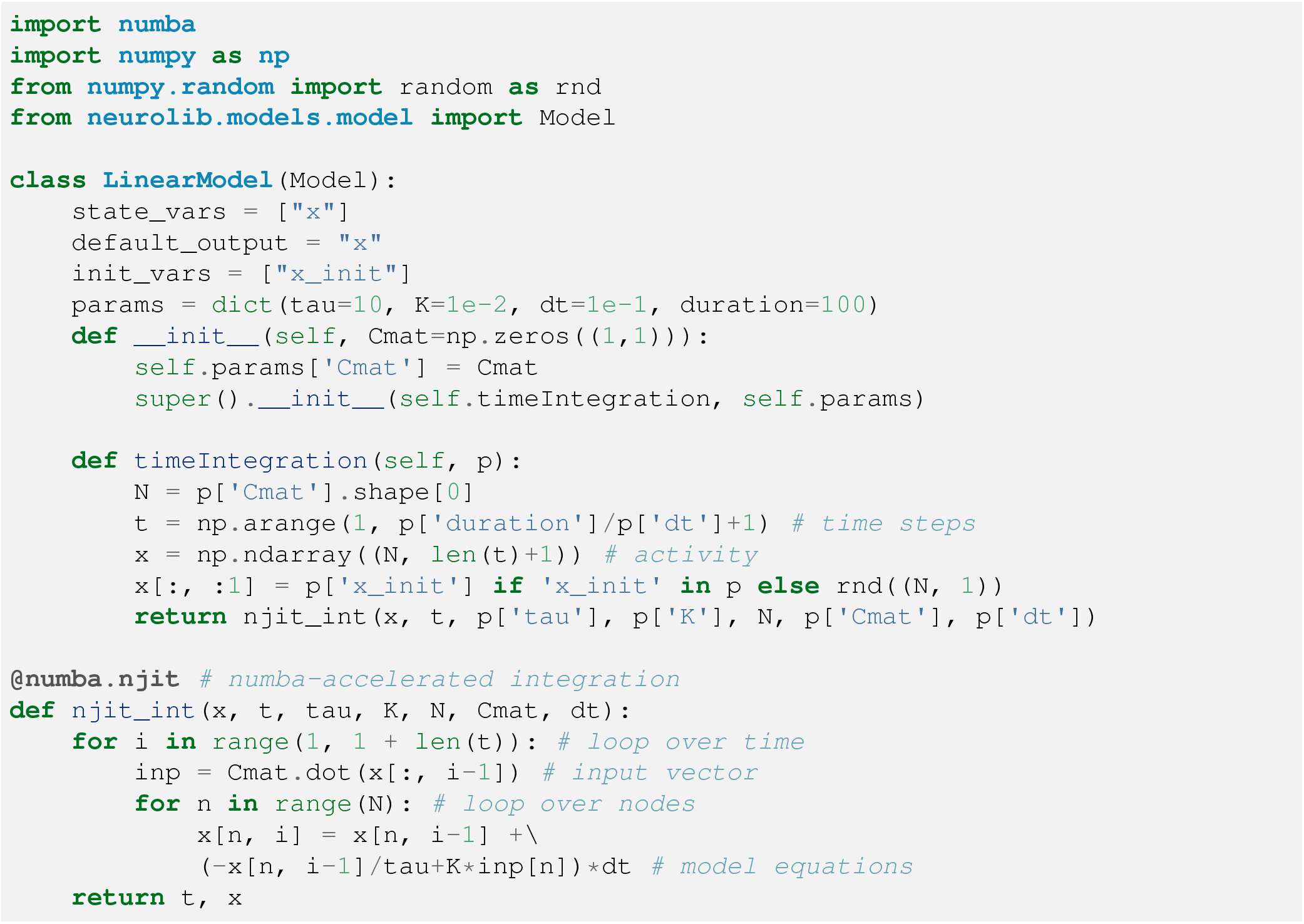
Custom model template. A linear model is implemented as a new class.

After the *i*-th chunk is integrated, only the last state vector of the system, **X**_i_(*T*_chunk_), is temporarily kept in memory, which in the case of a delayed system is a *n* × (*d_max_* + 1) matrix, with *n* being the number of state variables, and *d*_max_ being the number of time steps according to the maximum delay of the system. In the next step, all memory is cleared, and **x**_i_,(*T*_chunk_) is used as an initial state vector **x**_*i*+1_ (0) for the next chunk. If BOLD simulation is enabled, it will be integrated in parallel to the main integration and kept in memory, while the system’s past state variables, such as the firing rates, will be forgotten. After a long simulation is finished, the `output` attribute of the model will contain the long BOLD time series (e.g. 5 minutes) with a low sampling rate and the firing rates of the last simulated chunk (e.g. 10 seconds) with a high sampling rate.

### Parameter exploration

One of the main features of *neurolib* is its ability to perform parameter explorations in a unified way across models. Parameter explorations are useful for determining the behavior of a dynamical system when certain parameters are changed. For example, by measuring the minimum and maximum activity of a model given a parameter configuration, we can draw bifurcation diagrams that depict changes in the model’s dynamical state, for example transitions from constant activity to an oscillatory state. Figure 3 shows the bifurcation diagrams of the Hopf model, the Wilson-Cowan model, and the ALN model. Given these diagrams, we can choose the parameters of the system in order to produce a desired dynamical state. The time series in Fig. 3 show how the models behave at different points in the bifurcation diagrams. Currently, the exploration module of *neurolib* relies on *pypet*^45^ which manages the parallelization and the data storage of all simulations. The user can define the range of parameters that should be explored in a grid using the `ParameterSpace` class and pass it, together with the `model`, as an argument to the `BoxSearch` class (Listing 4). All simulated output will be automatically stored in an HDF5 file for later analysis.

**Listing 4.**
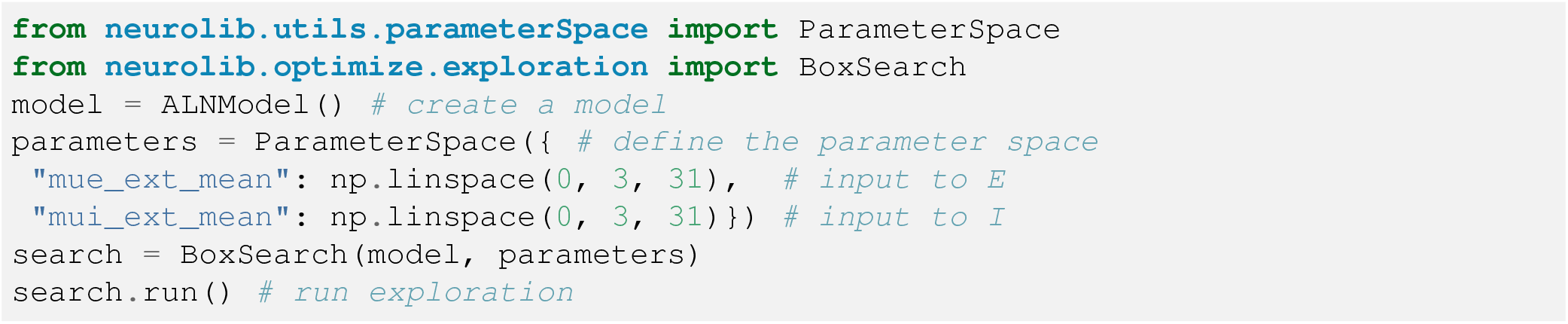
Parameter exploration. Setting up a parameter exploration of the ALN model using a grid search along the parameters 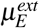 (`mue_ext_mean`) and 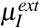 (`mui_ext_mean`) which represent the mean background inputs to the E and I subpopulations. These parameters represent the axes of the bifurcation diagram in Fig. 3 g.

Here, we used the *numpy*^46^ method `np.linspace` to define the parameters in a linear space between 0 and 3 in 31 steps. The `ParameterSpace` class then computes the Cartesian product of the parameters to produce a configuration for all parameter combinations. When the exploration is done, the results can be loaded from disk using `search.loadResults()`, which organizes all simulations and their outputs as a *pandas* DataFrame^47^ available as the attribute `dfResults`.

The result of this exploration is shown in Fig. 3 g as a two-dimensional state space diagram of a single node. A contour around states with a finite amplitude of the excitatory firing rate indicates bifurcations from states with constant firing rates to oscillatory states. Drawing bifurcation diagrams in terms of the external input parameters, as in Fig. 3 d and g is particularly useful, because it allows an interpretation of how the dynamics of an individual node in a brain network would change depending on the inputs from other brain areas. When the system is in the down-state, for example, enough excitatory external input from other brain areas would be able to push it over the bifurcation line into the oscillatory state.

It should be noted that, in the default implementation, all parameters are homogeneous across nodes, meaning that, in the example above, we change the external input currents to all brain regions at once. However, adjusting a model to work with heterogeneous parameters is easy and simply requires the use of vector-valued parameters instead of scalar ones. The numerical integration needs then to be adapted to use the elements of the parameter vector for each node accordingly.

The default behavior of `BoxSearch` is to simulate the model and store its outputs to disk. Alternatively, the user can pass the argument `evalFunction` which calls a separate function for every parameter combination instead. This enables the user to perform pre- and postprocessing steps for each simulation run. An example is shown in Listing 5.

**Listing 5.**
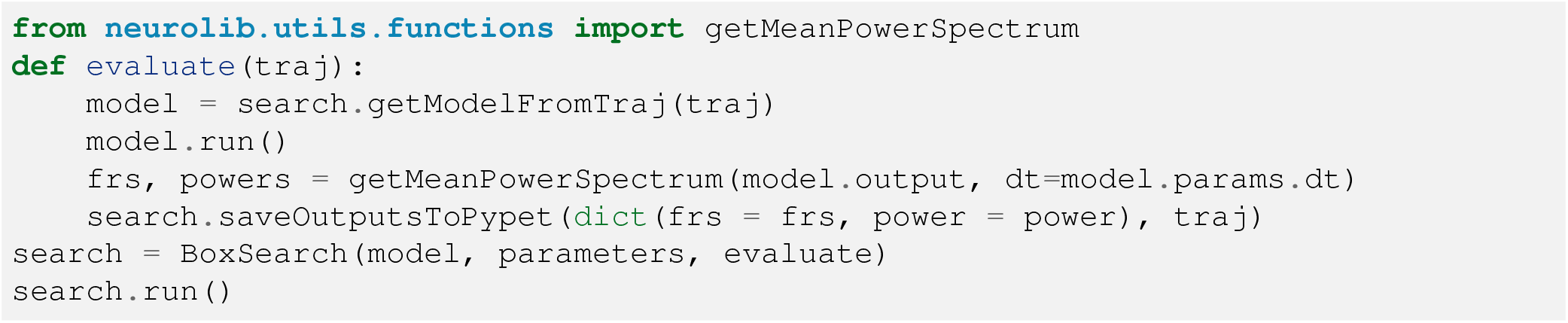
Parameter exploration with postprocessing. A custom function `evaluate()` is defined which runs the model and computes the power spectrum of its output using the builtin function `getMeanPowerSpectrum()`. The power spectrum is then saved to disk instead of the entire time series.

## Model optimization

Model optimization, in general, refers to a search for model parameters that maximize (or minimize) an objective function that depends on the output of a model, leading to a progression towards a predefined target or goal. In the case of whole-brain models, the goal is often to reproduce patterns from (i.e., fit the model to) empirical brain recordings. Optimizations can also be carried out for a single node. An example of this is finding local node parameters that produce an oscillation of a certain frequency, in which case the objective to minimize is the difference between the oscillation frequency of the model’s activity and a predefined target frequency, which could be based on, for example, empirical EEG or MEG recordings.

Whole-brain models are often fitted to empirical resting-state BOLD functional connectivity (FC) data from fMRI measurements. The goodness of fit to the empirical data is determined by computing the element-wise Pearson correlation coefficient of the simulated FC matrix and the empirical FC matrix. If the optimization is successful, the model produces similar spatial BOLD correlations as was measured in the empirical data. The FC Pearson correlation ranges between 0 and 1, where 1 means maximum similarity between simulated and empirical data.

To ensure that a model also produces temporal correlation patterns similar to fMRI recordings, the functional connectivity dynamics (FCD) matrix can also be fitted to empirical data. The FCD matrix measures the similarity of the temporal correlations between time-dependent FC matrices in a sliding window. The similarity between simulated and empirical matrices is usually determined by the Kolmogorov-Smirnoff (KS) distance^48^ between the distributions of the entries of both matrices. The KS distance ranges between 0 and 1, where 0 means maximum similarity.

In the whole-brain model, the relevant parameters that affect these correlations are typically the distance to a bifurcation line that separates the steady-state from an oscillatory state (see Figs. 3 a, d, and g), the global coupling strength *K* that determines how strongly all brain regions are coupled with each other, and other parameters, such as the signal transmission speed or the strength of the external noise.

### Evolutionary algorithms

*neurolib* supports model optimization through evolutionary algorithms built using the evolutionary algorithm framework *DEAP*^49^. Evolutionary algorithms are stochastic optimization methods that are inspired by the mechanisms of biological evolution and natural selection. Multi-objective optimization methods, such as the NSGA-II algorithm^50^, are crucial in a setting in which a model is fit to multiple independent targets. These could be features from fMRI recordings, such as FC and FCD matrices, or from other data modalities such as EEG. In a multi-objective setting, not one single solution but a set of solutions can be considered optimal, called the Pareto front, which refers to the set of solutions that cannot be improved in any one of the objectives without diminishing its performance in another one. These solutions are also called non-dominated.

In the evolutionary framework, a single simulation run is called an individual. Its particular set of parameters are called its genes and are represented as a vector with one element for each free parameter that should be optimized. A set of individuals is called a population. For every evolutionary round, also called a generation, the fitness of every new individual is evaluated by simulating the individual and computing the similarity of its output to the empirical data, as shown in Listing 6. In our example, this results in a two-dimensional fitness vector with FC and FCD fits.

A schematic of a general evolutionary algorithm is shown in Fig. 4. The optimization can be separated into an initialization phase and the main evolutionary loop. In the initialization phase, all individuals are uniformly sampled from the parameter space and are evaluated for their fitness. The optimization then enters the second phase in which the population is reduced to a subset of size *N*_pop_ from which parents are selected to generate offspring for the next generation. These offspring are mutated, added to the total population, and the procedure is repeated until a stopping condition is reached, such as a maximum number of generations.

**Figure 4.**
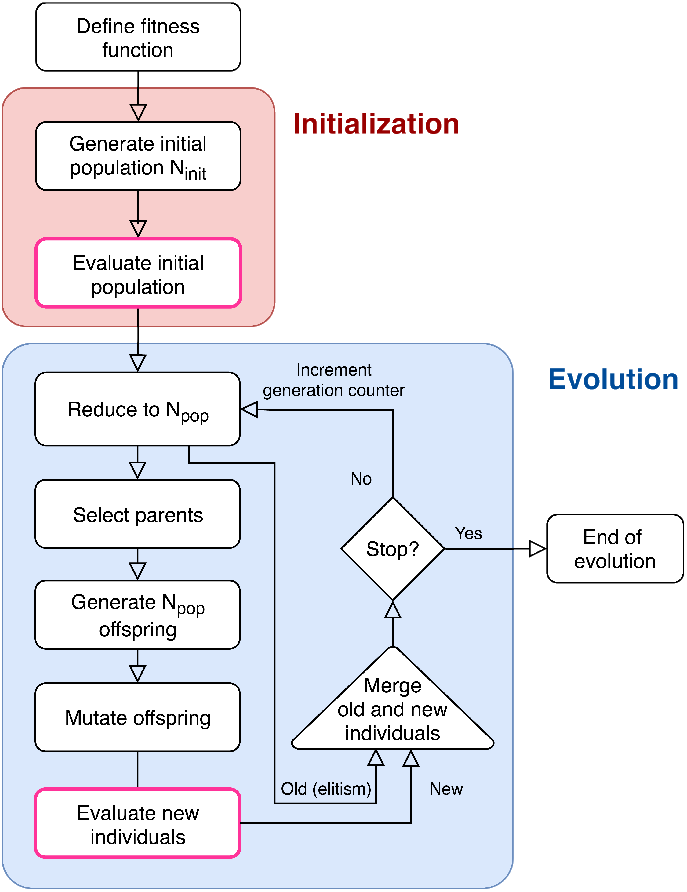
Schematic of the evolutionary algorithm. The optimization consists of two phases, the initialization phase (red) in which the parameter space is uniformly sampled, and the main evolutionary loop (blue). N_init_ is the size of the initial population, N_pop_ is the size of the population during the evolution.

**Listing 6.**
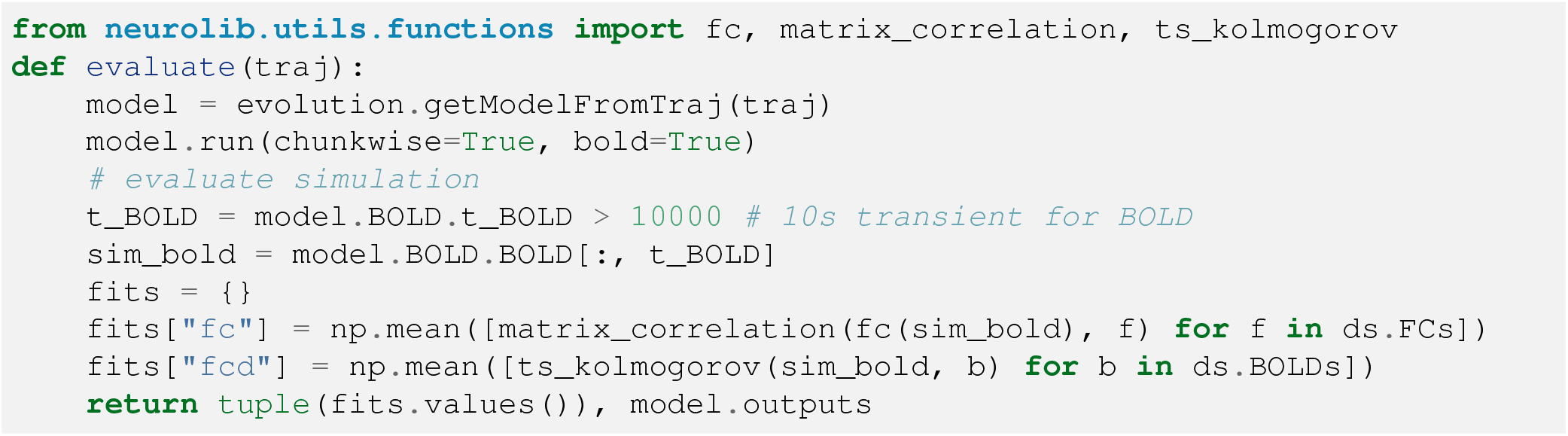
Evaluation of model fitness during evolution. After loading and simulating the model, the fitness is determined by comparing its BOLD output to the empirical fMRI data. The simulated BOLD time series is measured after a 10 second transient period. The functional connectivity (FC) is computed using the function `fc()`. The fit to empirical FC matrices is computed using the function `matrix_correlation()`. The fit to empirical functional connectivity dynamics (FCD) matrices is calculated using the function `ts_kolmogorov()`. Fits are determined for each subject of the dataset ds and averaged across all subjects.

**Listing 7.**
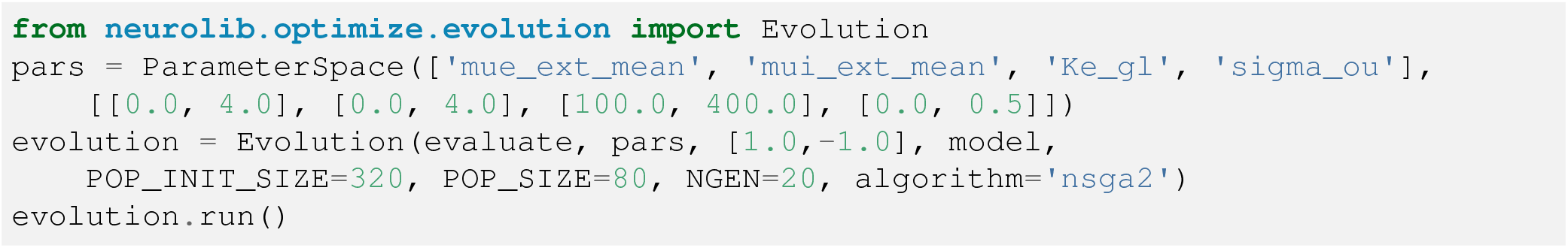
Evolutionary optimization. Multi-objective optimization using the NSGA-II algorithm. Four parameters are optimized, namely the background inputs 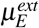 (`mue_ext_mean`) and 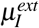 (`mui_ext_mean`), the global coupling strength *K* (`Ke_gl`) and the noise strength σ_ou_ (`sigma_ou`, see Eq. 2). The first argument of `Evolution` refers to the `evluate` function in Listing 6, and the third argument indicates that the fitness vector is two-dimensional, and that the first value (FC correlation) will be maximized (+1), and the second one (FCD distance) will be minimized (−1). The initial population size of the evolution is set to 320, the main population size to 80, and the number of generations is set to 20.

As an alternative to the NSGA-II algorithm, which is particularly useful for multi-objective optimization settings, an adaptive evolutionary algorithm^51^ is also supported. In the adaptive algorithm, the mutation step size, which is analogous to a learning rate, is also learned during the evolution by treating it as an additional gene during the optimization. This algorithm has been successfully used to optimize whole-brain models with up to 8 free parameters, however, it is not guaranteed to produce satisfactory results in a multi-objective optimization setting. All steps in the evolutionary algorithm can be modified by implementing custom operators or by using the ones available in *DEAP*.

## Example: evolutionary model optimization

In the following, we show how an evolutionary optimization is set up in *neurolib* using the NSGA-II algorithm. We use the brain network model from Listing 2 and fit the BOLD output of the model to the empirical BOLD data to capture its spatiotemporal properties, as described above. In Listing 7, we specify the parameters to optimize and their respective boundaries, and define a weight vector that determines whether each fitness value is to be maximized (+1) or minimized (−1). In our case, we want to maximize the first measure, which is the FC correlation, and minimize the second, which is the FCD distance. The size of the initial population *N*_init_, the ongoing population *N*_pop_ size, and the number of generations *N*_gen_ largely determine the time for the optimization to complete.

We visualize the increasing score of the population in Fig. 5 a, defined as the weighted sum of all objectives, as the evolution progresses. As expected, we find all good fits (with FC correlation > 0.35 and FCD distance < 0.5) close to the bifurcation line between the down-state and the limit cycle, shown in Fig. 5 b. Note that while we have optimized 4 parameters simultaneously, only the input current parameters to E and I are shown, which correspond to the axis of the bifurcation diagram in Fig. 3 g. In another example, we also included EEG power spectra in our optimization procedure to produce a model with an appropriate spectral density. Here, we used EEG data from sleep recordings during the sleep stage N3^20,52^, or slow-wave sleep, in which sleep slow oscillations are prevalent. In order to assess the fit to the power spectrum, we computed the mean of the power spectra of the excitatory firing rate of all nodes during the last 60 seconds of the simulation using the function `getMeanPowerSpectrum()` which uses the implementation of Welch’s method^53^ *scipy.signal.welch* in SciPy^37^ with a rolling Hanning window of length 10s. The same method was applied to the channel-wise EEG data to first compute subject-wise average power spectra, and then average all subject-wise spectra to a single empirical power spectrum. To assess the similarity of the simulated and the empirical data, we computed the Pearson correlation between both power spectra in a range between 0 and 40 Hz.

**Figure 5.**
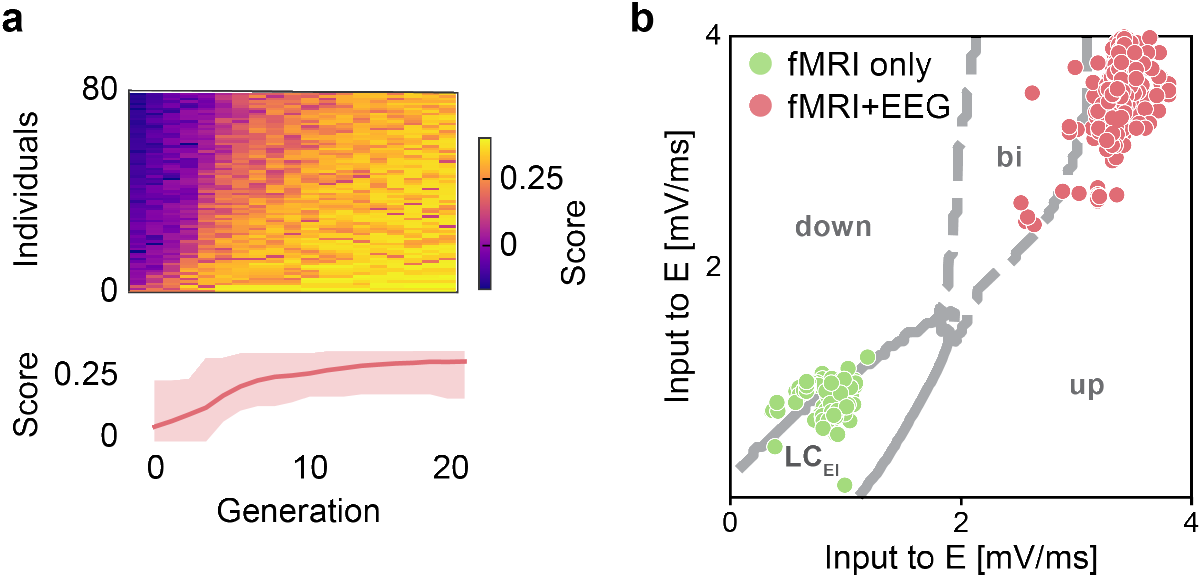
Optimization result. **(a)** Improvement of the fitness over all generations with the color indicating the score per individual (top row) and averaged across the population (bottom row) with the minimum and maximum range in the shaded area. The fitness score is a weighted sum of all individual fitness values, i.e. higher values are better. **(b)** Evolution results of the optimal parameters for the input currents to the E and I subpopulations depicted in the bifurcation diagram of the ALN model. Green dots indicate optimal parameters for fits to fMRI data only (progression shown in a), red dots show fits to fMRI+EEG simultaneously. The resulting fits of the fMRI+EEG case are shown in Fig. 6. In the fMRI-only case, no spike frequency adaptation mechanism was used. In the fMRI+EEG case, the adaptation strength *b* and timescale *τ_A_* were allowed to vary.

In order for the model to produce slow oscillations that fit the data, we included the spike-frequency adaptation strength parameter *b* and the adaptation time scale *τ_A_* in our optimization, culminating in a total of six free parameters. In a single node, adaptation-induced oscillations can have a frequency between roughly 0.5 and 5 Hz^24^. Figure 5 b shows the location of all good fits in the bifurcation diagram (FC and FCD thresholds as above, EEG power spectrum correlation > 0.7). All fits are close to where the bistable regime is in the case without adaptation, where the activity of an E-I system can slowly oscillate between up- and down-states if the adaptation mechanism is strong enough^24,34,54^.

Figures 6 a-c show the simulated and empirical FC and FCD matrices, as well as the power spectra of a randomly chosen fit of the fMRI+EEG optimization from Fig. 5 b. The empirical FC and FCD matrices are shown for one subject only. The correlation between simulated and empirical FC matrices was 0.55 averaged across all subjects with the best subject reaching 0.70. The KS distance of the distribution of FCD matrix entries averaged across all subjects was 0.28 with the best subject reaching 0.07. The correlation coefficient between the power spectra of the simulated firing rate EEG was 0.86.

**Figure 6.**
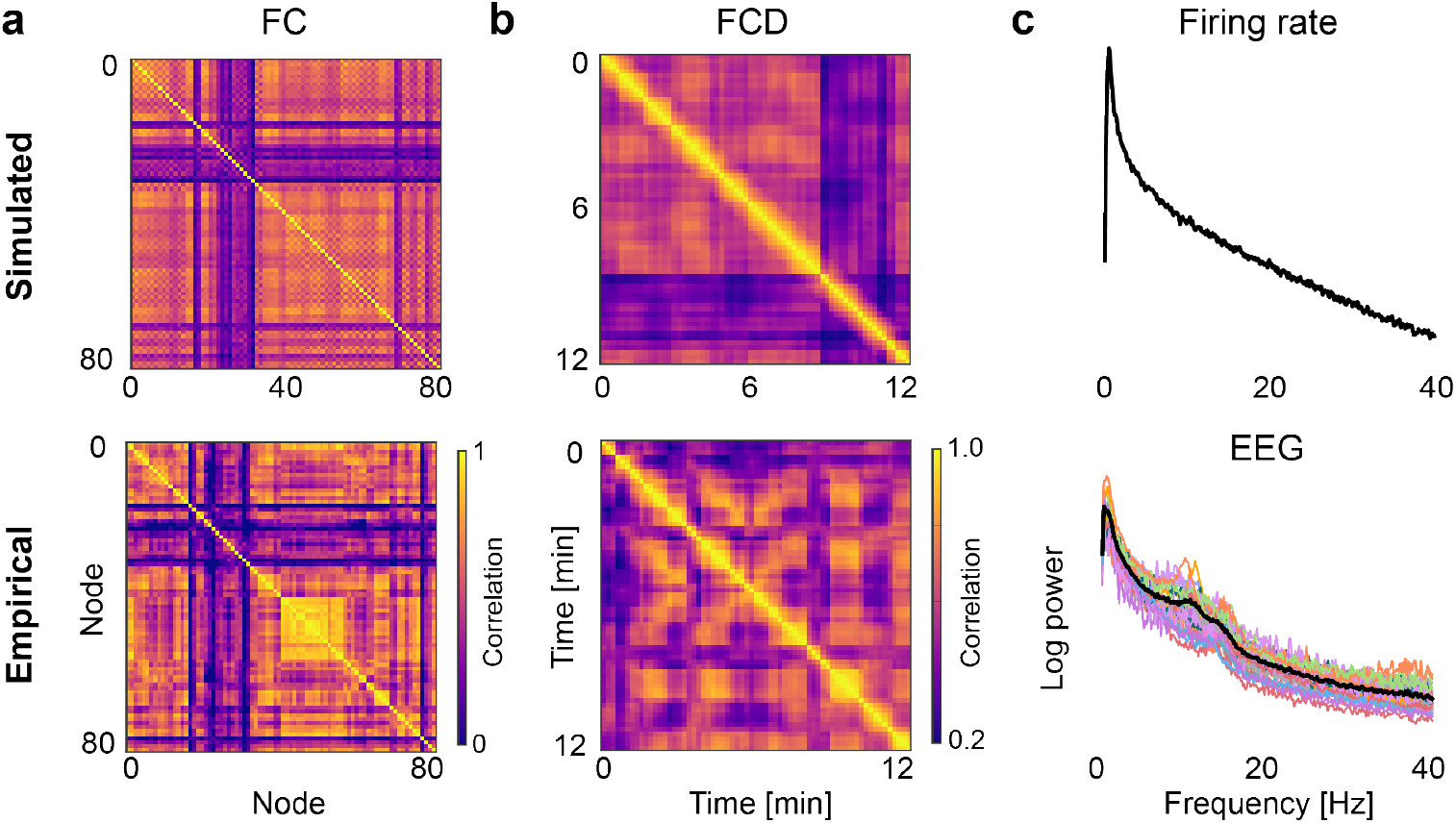
Optimal fits to empirical data. **(a)** Simulated (top panel) and empirical (bottom panel) FC matrices, **(b)** FCD matrices, and **(c)** the average power spectrum of the simulated firing rate and empirical EEG data during sleep stage N3. The spectra of the individual subjects is shown in color, the average spectrum is shown in black.

## Summary and discussion

In this paper, we introduced *neurolib*, a Python library for simulating whole-brain networks using coupled neural mass models. We demonstrated how to simulate a single neural mass model, how *neurolib* handles empirical data from fMRI and DTI measurements, and how to simulate a whole-brain network. A set of neural mass models that are part of the library was presented, as well as how to implement a custom neural mass model.

We demonstrated how to conduct parameter explorations with *neurolib* which can be used to characterize the dynamical landscape of a model. Lastly, we presented how the multi-objective evolutionary optimization algorithm in *neurolib* can be employed to fit a whole-brain model to functional empirical data from fMRI and EEG.

### Existing software

A well-established alternative to *neurolib* is *The Virtual Brain (TVB*)^55,56^ which is an easy-to-use platform for running brain network simulations. *TVB* can load structural connectivity data, has a long list of implemented models for simulating brain regions, and allows users to set up monitors to record activity. *TVB* can also simulate BOLD signals and various other forward models, such as simulated electroencephalography (EEG), magnetoencephalography (MEG) and local field potentials (LFP). Many of the features of *TVB* can be accessed and configured using a graphical user interface (GUI), however more complex use cases, such as fitting a model to empirical data, or further analyzing model outputs, need to be managed outside of the graphical environment.

In contrast, *neurolib* does not have a GUI and encourages users with programming experience to modify the code of the framework itself to suit their individual use case, to implement their own models, and to use their own datasets to run large numerical experiments. *neurolib* also offers parameter exploration and model optimization capabilities. The simple and efficient architecture of *neurolib* allows for fast prototyping of custom models. Models can be implemented directly in Python and the numerical integration can be accelerated using the just-in-time compiler *numba.*

Although *TVB* also uses accelerated *numba* code, only the calculation of the derivatives of the models are accelerated but not the integration itself. In *neurolib*, the entire numerical integration, including the loops across all nodes of the brain network, are accelerated, resulting in a simulation speed that is 8-24x faster compared to *TVB* (see Figure 7).

**Figure 7.**
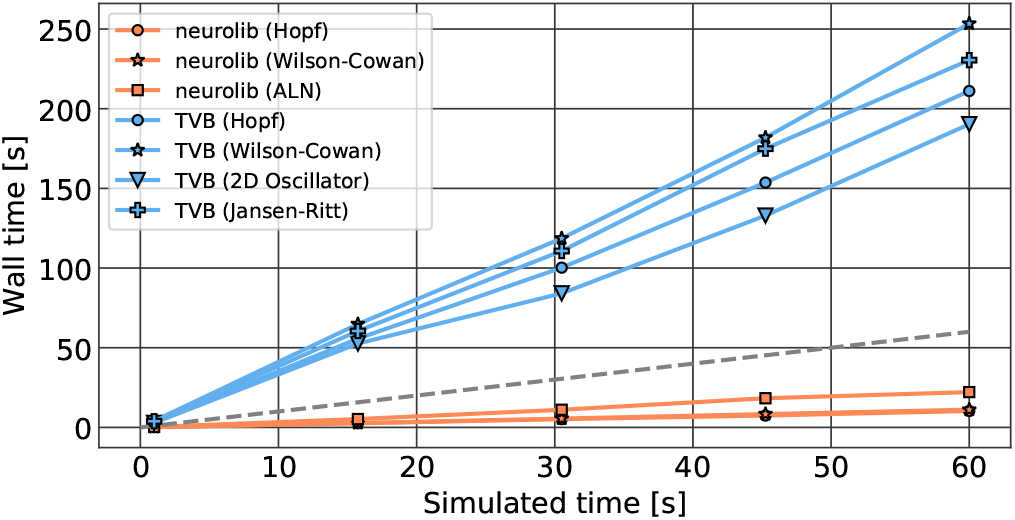
Performance comparison. Wall time (real time) plotted against the simulated time in *neurolib* (orange) and *The Virtual Brain (TVB)* (blue) for different models. The dashed gray line indicates the identity line at which simulated time is equal to wall time. At each data point, a whole-brain model with 76 brain regions was simulated. Before each measurement, the simulation was run once to avoid additional waiting times due to precompilation of the code. Performance was measured on a MacBook Pro 2019 with a 2.8 GHz Quad-Core Intel Core i7 CPU.

### Future development

Current work on *neurolib* focuses on a new feature called *multimodel*, which achieves heterogeneous brain modeling in which more than one type of neural mass model is used to simulate a brain network. This should not be confused with using the same model with different parameters for each of the *N* brain areas, a functionality that can be easily implemented within the existing framework by using *N*-dimensional vector-valued parameters instead of scalar ones. Heterogeneous brain modeling requires the user to define which output variables of each model should be coupled to which input variables of another. Keeping tabs on which data needs to be exchanged between brain areas and synchronizing different parts of the simulation, while ensuring that computational efficiency does not suffer too much, has proven to be a great challenge in this endeavor. The goal of this will be that, in the future, we will be able to run whole-brain simulations using specialized models for different parts of the brain, such as by combining a cortical model (like the ALN model) with models of thalamic or hippocampal neural populations.

Another goal of the development efforts is to support more sophisticated forward models like the ones used in TVB. This includes making use of lead-field matrices to simulate an EEG/MEG signal that is more spatially accurate in sensor space, making comparisons to real recordings more faithful than by simply analyzing neural activity in source space.

## Conclusion

The primary development philosophy of *neurolib* is to build a framework that is lightweight and easily extensible. Future work will also include the implementation and support for more neural mass models. Since *neurolib* is open source software, we welcome contributions from the computational neuroscience community. Lastly, our main focus in developing *neurolib* is the computational efficiency with which simulations, explorations, and optimizations can be executed. We believe that this not only has the potential to save valuable time, but allows researchers to pursue ideas and conduct numerical experiments that would otherwise be only achievable with access to a large computing infrastructure.

## Code availability and documentation

All of our code including examples and documentation can be accessed on our public Github repository which can be found at https://github.com/neurolib-dev/neurolib. The documentation including more examples of how to use *neurolib* can be found at https://neurolib-dev.github.io/.

## Acknowledgments

This work was funded by the Deutsche Forschungsgemeinschaft (DFG, German Research Foundation) – Project number 327654276 - SFB 1315 and by the Operational Programme Research, Development and Education, Ministry of Education, Youth and Sport of the Czech Republic (co-funded by the EU) - project no. CZ.02.2.69/0.0/0.0/19_074/0016209

## Notes

### Competing Interest Statement

The authors have declared no competing interest.

